# Sphingosine-1-phosphate receptor modulators resensitize FLT3-ITD acute myeloid leukemia cells with *NRAS* mutations to FLT3 inhibitors

**DOI:** 10.1101/2025.11.21.689510

**Authors:** Aditi Chatterjee, Moaath K. Mustafa Ali, Christopher M. Bailey, Yuchen Liu, Donald Small, Catherine C. Smith, Elie Traer, Yin Wang, Giovannino Silvestri, Maria R. Baer

**Author notes:** **Correspondence to:** Maria R. Baer, MD, University of Maryland Greenebaum Comprehensive Cancer Center, 22 South Greene Street, Baltimore, MD 21201.

## Abstract

FLT3 inhibitor efficacy in AML with FLT3-ITD is short-lived, frequently due to new mutations, most commonly in *NRAS*. Sphingosine kinase 1 (SPHK1), which phosphorylates sphingosine to generate sphingosine-1-phosphate (S1P), is upregulated and localized to the plasma membrane in *RAS*-mutated cells. We studied S1P and FLT3 co-targeting to overcome FLT3 inhibitor resistance in *NRAS*-mutated FLT3-ITD AML cells. *NRAS*-mutated FLT3-ITD AML cell lines and patient blasts were treated with FLT3 inhibitors and/or S1P receptor (S1PR) modulators. FLT3 inhibitor sensitivity was assessed by immunoblotting, cytotoxicity and apoptosis assays. Co-treatment was also assessed *in vivo* in an orthotopic mouse model. Downstream RAS and SPHK1 effectors were measured by immunoblotting and qRT-PCR. The S1PR modulators fingolimod (FTY720) and mocravimod (KRP-203) resensitized FLT3-ITD-expressing MOLM-14 and MV4-11 human AML cells with G12D, G12S, Q61K or Q61H, but not G12C, and patient blasts with G13D or G13V *NRAS* mutations to FLT3 inhibitors. Moreover, FTY720 co-treatment resensitized G12D *NRAS*-mutated M14(R)701 cells to gilteritinib *in vivo.* Co-treatment inactivated ERK, transcriptionally downregulated SPHK1, and inactivated downstream AKT, p70S6K and BAD, with inactivation abrogated by constitutive SPHK1 expression. The clinically applicable S1PR modulators fingolimod and mocravimod resensitize *NRAS*-mutated FLT3-ITD AML cells to FLT3 inhibitors, supporting potential clinical efficacy of these combinations.

## Introduction

*FMS*-like tyrosine kinase 3 internal tandem duplication (FLT3-ITD) is present in acute myeloid leukemia (AML) cells in 25% of patients, associated with rapid relapse after treatment response and poor outcomes (1,2). Wild-type FLT3 signals through the PI3 kinase (PI3K)-AKT-mTOR and Ras-Raf-MEK-ERK pathways upon binding of FLT3 ligand (3), and FLT3-ITD signals constitutively through these two pathways, as well as, aberrantly, though signal transducer and activator of transcription (STAT) 5, promoting AML blast proliferation and resistance to apoptosis (3). FLT3 tyrosine kinase inhibitors improve outcomes in FT3-ITD AML (4–6), but efficacy is short-lived (5), with onset of resistance often associated with new mutations (7).

*NRAS* is the most commonly mutated gene in FLT3-ITD AML with acquired FLT3 inhibitor resistance (8–13). The RAS family proteins NRAS, HRAS and KRAS encode structurally similar small GTPase signal transduction proteins that are activated downstream of receptor tyrosine kinases, including FLT3, following ligand binding (3). Point mutations altering RAS protein codons G12, G13 or Q61 impair GTPase activity, constitutively activating RAS proteins and Ras-Raf-MEK-ERK signaling (14). *NRAS* mutations reported to date in FLT3-ITD AML include G12D, G12S, G12C, Q61K, Q61R, Q61K, Q61H, Q61R, G13D, G13C and G13R (10).

Co-targeting other proteins altering FLT3-ITD signaling may enhance FLT3 inhibitor efficacy and prevent or overcome resistance (15). Sphingosine kinase 1 (SPHK1), which catalyzes phosphorylation of the lipid signaling molecule sphingosine to generate sphingosine-1-phosphate (S1P), is upregulated and localized to the plasma membrane in an ERK-dependent manner in *RAS*-mutated cells (16,17). We hypothesized that targeting S1P signaling in conjunction with inhibiting FLT3 could overcome FT3 inhibitor resistance in *NRAS*-mutated FLT3-ITD AML cells.

## Materials and methods

### Cell lines

MOLM-14 and MV4-11 human AML cells, with heterozygous and homozygous FLT3-ITD, respectively, were obtained and cultured as described (18). M14(R)701 cells, with a G12D activating *NRAS* mutation, were generated by culturing MOLM-14 cells with lestaurtinib (CEP-701) (8). M14(R)701 cells were cultured in RPMI 1640 medium with 10% fetal bovine serum, 1% penicillin-streptomycin and 10 nM CEP-701, but without CEP-701 for at least 2 days before initiating experiments (8). Quizartinib-resistant MOLM-14(QS)-NRASQ61K and MOLM-14(QS)-NRASG12C cells, with *NRAS* Q61K and G12C mutations, respectively, were generated by culturing MOLM-14 cells in medium containing quizartinib in escalating concentrations (0.5 to 20 nM), (10). Gilteritinib-resistant MOLM-14 NRAS G12S cells were generated by culturing MOLM-14 cells with 100 nM gilteritinib and 10 ng/mL fibroblast growth factor 2 (FGF2), and gilteritinib-resistant MV4-11 NRAS Q61H cells by culturing MV4-11 cells with 100 nM gilteritinib and 10 ng/mL FLT3 ligand (13).

### Lentiviral infection

M14(R)701 cells were infected with pLenti-GIII-CMV vector containing the SPHK1 gene [SPHK1 Lentiviral Vector (Human) (CMV) (pLenti-GIII-CMV, Retired Cat. No.: 45312062, Applied Biological Materials, Richmond, Canada]. pLenti X1 Puro empty vector (plasmid 20953, Addgene, Lewisville, TX) served as the control. Lentiviral particles were generated through transient calcium phosphate transfection using the ProFection Mammalian Transfection System (E1200, Promega, Madison, WI), applied to HEK293T cells (CRL-3216, ATCC, Manassas, VA). Viral supernatant collected 24 and 48 hours after transfection was concentrated by centrifugation with Viro-Peg 100 Lentivirus Concentrator (CHM00B952, Thomas Scientific, Swedesboro, NJ). M14(R)701 cells underwent three rounds of spinoculation in the presence of 4 µg/mL polybrene (TR-1003-G, Millipore Sigma, Burlington, MA). Puromycin (A1113803Gibco, Grand Island, NY) selection was initiated 48 hours after transduction.

### Patient samples

Blood samples were obtained from patients with FLT3-ITD AML and *NRAS* mutations with hyperleukocytosis and peripheral blasts (Supplementary Table S1) on a University of Maryland School of Medicine Institutional Review Board-approved tissue procurement protocol, following written informed consent. Studies were conducted in accordance with the Declaration of Helsinki. Mononuclear cells were isolated by density centrifugation over Ficoll-Paque (GE17-1440-02, Millipore Sigma).

### Materials

FLT3 inhibitors included lestaurtinib (CEP-701; HY-50867, MedChem Express, Monmouth Junction, NJ) and gilteritinib (ASP 2215; S7754) (Type I), and quizartinib (AC220, S1526) (Type II) (Selleck Chemicals, Houston, TX). Sphingosine-1-phosphate receptor (S1PR) modulators included fingolimod (FTY720; SML0700, Sigma-Aldrich, St. Louis, MO) and mocravimod (KRP-203; HY-13660, MedChemExpress). Other drugs included the protein phosphatase 2A (PP2A)-activating drug (PAD) DT-061 (HY-112929, MedChemExpress), and the protein translation inhibitor cycloheximide (CHX; C7698) and proteasome inhibitor carbobenzoxy-L-leucyl-L-leucyl-L-leucinal (MG-132; 474790), both from Sigma-Aldrich.

### Cytotoxicity assays

Drug cytotoxicity was measured using the WST-1 assay (18). Briefly, cells plated at 10,000/well in triplicate were treated for 48 hours with FLT3 inhibitors at increasing concentrations alone and in combination with FTY720 or DT-061 at fixed concentrations. Absorbance was read after incubation with WST-1 Cell Proliferation Reagent (11644807001, MilliporeSigma) for 2 hours. Cytotoxicity of single drugs and combinations was analyzed using Prism 10 (GraphPad Software, San Diego, CA).

### Drug Combinations

Cells seeded on 96-well plates were treated in triplicate with drugs at diverse concentrations alone and in combinations, and cytotoxicity was measured as above (19). Drug combination effects were analyzed using the Chou-Talalay method, with CompuSyn software (CompuSyn, Paramus, NJ) (19). Combination index less than, equal to, and greater than 1.0 indicated synergy, additivity, and antagonism, respectively (20).

### Measurement of apoptosis

Cells were stained with Annexin V (556419) and propidium iodide (556463) (BD Biosciences, Franklin Lakes, NJ) and analyzed on a FACS Canto II flow cytometer (BD Biosciences) with FlowJo v10.9 software (BD Biosciences) (19). Percent total annexin V+/PI− and annexin V+/PI+ cells was determined.

### in vivo study

M14(R)701 cells were transfected with luciferase and green fluorescent protein (GFP). Exponentially growing cells (0.5×10^6^) were injected into the lateral tail veins of restrained female NOD-Rag1null IL2rgnull, NOD rag gamma (NRG) mice (6-8 weeks old). Engraftment was assessed using an IVIS Lumina LT Series III imaging system (PerkinElmer, Shelton, CT) after intraperitoneal injection of 150 mg/kg D-luciferin (50227, MilliporeSigma). Mice were sorted into 4 groups of 5 with equal mean signal intensity. Treatment was initiated on day 12 post-injection with gilteritinib 7.5 mg/kg (19) in 5% DMSO, 40% polyethylene glycol 300 (PEG 300), 5% polysorbate 80 (Tween 80) and 50% water by oral gavage 5 days/week or vehicle control, and/or FTY720 10 mg/kg (21) dissolved in DMSO and prepared as 250 µg/200 µl 0.9% saline solution and injected intraperitoneally 5 days/week, or vehicle control. Weights were recorded before each treatment. Leukemia burden was assessed weekly by non-invasive luciferin imaging. Bioluminescent image data were analyzed with Living Image software (128113, Revvity, Waltham, MA). Endpoints were 20% body weight loss, hind limb paralysis or lack of mobility to eat/drink. The University of Maryland Institutional Animal Care and Use Committee approved the study.

### PP2A phosphatase assay

PP2A phosphatase activity was measured using the PP2A Immunoprecipitation Phosphatase Assay Kit (17-313, MilliporeSigma) (21). Briefly, protein lysates (50 μg) in 100 μL 20 mM HEPES (pH 7.0), 100 mM NaCl, 5 μg PP2Ac antibody and 25 μL protein A-agarose were added to 400 μL 50 mmol/L Tris (pH 7.0)/100 mM CaCl_2_. Immunoprecipitation was carried out at 4°C for 2 hours. Immunoprecipitates were used in the phosphatase reaction.

### Immunoblotting

Lysate (18,19) immunoblots were probed with antibodies to p-STAT5 (Y694) (9351), STAT5 (94205), p-p44/42 MAPK (Erk1/2; T202/Y204) (9101), p44/42 MAPK (Erk1/2) (9107), p-Akt (S473) (T308) (4060) (4056), Akt (9272), p-GSK-3α/β (S21/9) (8566), GSK-3β (9832), SPHK1 (12071), p-p70 S6K (T389) (9206), p70 S6K (9202), p-BAD (S136) (4366), BAD (9239) (Cell Signaling Technology, Danvers, MA), and vinculin (V9264) (Invitrogen, Waltham, MA) at 1:1000 dilution overnight at 4°C, then with horseradish peroxidase-conjugated secondary antibodies for one hour at room temperature. Band intensities quantified by densitometry (Image J, NIH) were normalized to vinculin.

### Protein turnover

Cells were treated with 100 µg/mL cycloheximide (CHX; C7698, Millipore Sigma) for 60 minutes to block new protein translation before adding FLT3 inhibitor and/or S1PR modulator, or DMSO control. Protein expression was measured at serial time points by immunoblotting. Band intensities quantified by densitometry were normalized to vinculin and compared to Time 0 (pre-treatment) levels. The line of best fit was used to determine 50% protein turnover time points (19).

### Quantitative real time polymerase chain reaction (qRT-PCR)

RNA isolated from cell lines in triplicate using RNeasy Plus (74134, Qiagen, Germantown, MD) was measured using a NanoDrop™ Lite Spectrophotometer (Thermo Fisher, Waltham, MA). RNA (500 ng) from each sample was reverse-transcribed using the SuperScript^TM^ IV First-Strand Synthesis System (18091050, Thermo Fisher). Each gene was amplified in triplicate using PowerUp SYBR Green Master Mix (4367659, Applied Biosystems, Waltham, MA) in the CFX Connect Real-Time PCR Detection System (Bio-Rad, Hercules, CA). Primers are listed in Supplementary Table S2. mRNA expression levels were determined using the ΔCt method for relative quantification. mRNA levels were normalized to 18S mRNA levels at serial time points and compared to Time 0 (pre-treatment) levels, defined as 1.

### Statistical analysis

Statistical analysis was performed on three independent experiments by unpaired t-test, using Prism 10 (GraphPad, San Diego, CA). In the *in vivo* model, photon intensity in mice treated with FTY720 and gilteritinib combination versus gilteritinib alone was compared by two-way ANOVA with Sidak’s multiple comparison test, and survival was compared by Kaplan-Meier analysis (18,19).

## Results

### *NRAS*-mutated MOLM-14 cells are resistant to gilteritinib and quizartinib

Parental and *NRAS* G12D-, G12C-, G12S- and Q61K-mutated MOLM-14 cells and parental and NRAS Q61H-mutated MV4-11 cells were cultured with gilteritinib or quizartinib and with FTY720 or KRP-203 in increasing concentrations. All *NRAS*-mutated cells were resistant to both FLT3 inhibitors, with 4- to >4,000-fold higher IC_50_s than parental cells, but parental and *NRAS*-mutated cells had similar FTY720 and KRP-203 IC_50_s (Supplementary Table S3).

### S1PR modulators FTY720 and KRP-203 resensitize M14(R)701 cells to FLT3 inhibitors

*NRAS* G12D-mutated M14(R)701 cells were cultured with gilteritinib or quizartinib at increasing concentrations with and without FTY720 at 2.5 µM, a concentration that is not toxic to normal primary hematopoietic progenitors (21). Co-treatment with FTY720 sensitized M14(R)701 cells to both gilteritinib and quizartinib (Figure 1A). Similarly, FTY720 combined with gilteritinib or quizartinib at different combinations of concentrations produced synergistic cytotoxicity in M14(R)701 cells by Chou-Talalay analysis (Figure 1B).

**Figure 1.**
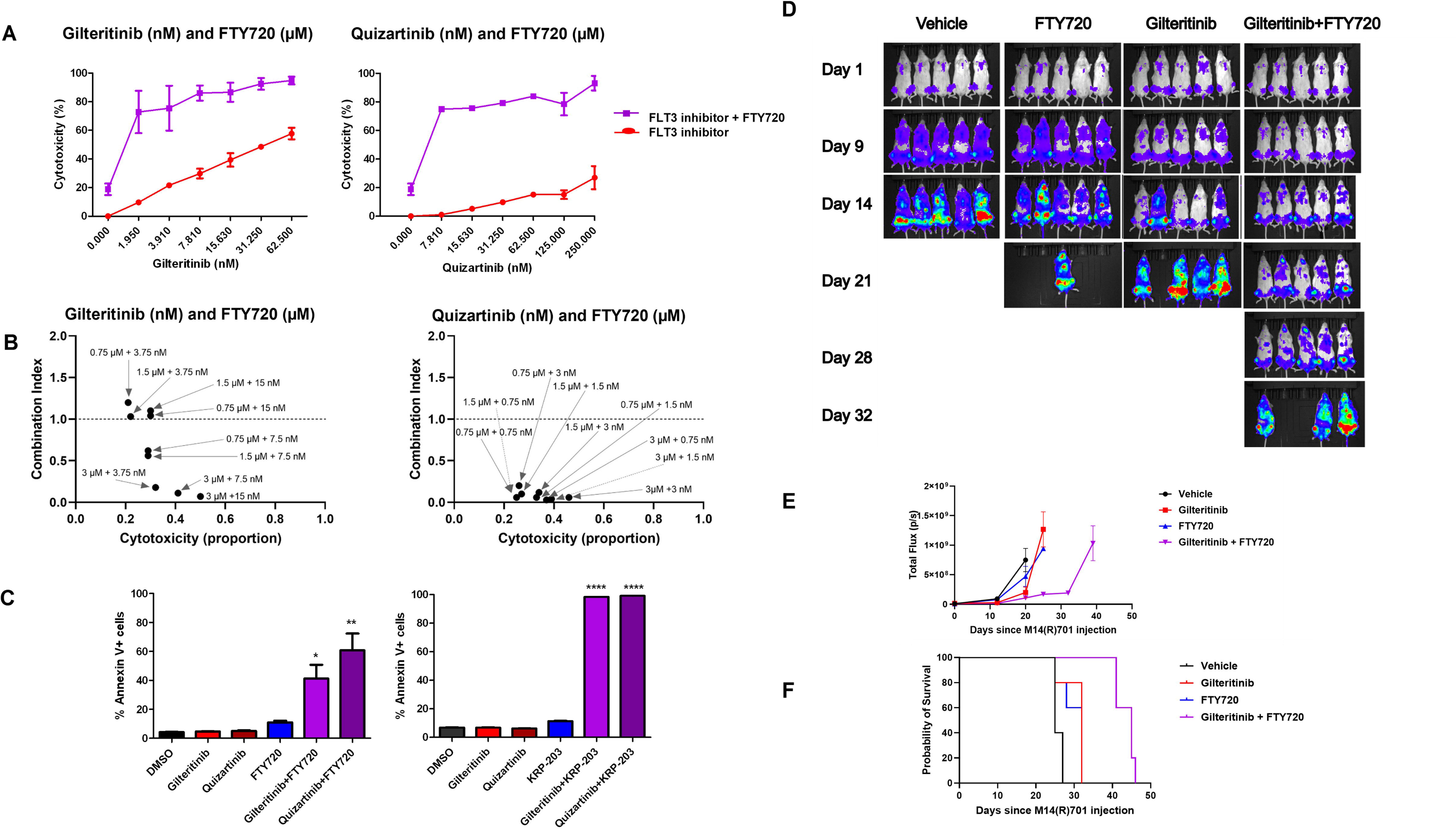
S1PR modulators resensitize M14(R)701 cells to FLT3 inhibitors *in vitro* and *in vivo*. **A.** M14(R)701 cells were treated with the FLT3 inhibitors gilteritinib and quizartinib at increasing concentrations alone and in combination with the S1PR modulator FTY720 (2.5 μM) for 48 hours. Cytotoxicity was measured with the WST1 assay. **B.** M14(R)701 cells seeded at 10,000 cells/well in 96-well plates were treated for 48 hours with gilteritinib or quizartinib and/or FTY720 as single drugs and in combinations at the concentrations shown, in triplicate. Cytotoxicity was measured with the WST-1 assay, and drug combination effects by the Chou-Talalay method. Synergy was defined by combination index values < 1.0. **C.** M14(R)701 were treated with gilteritinib (10 nM), quizartinib (1 nM), FTY720 (2.5 μM) or KRP-203 (5 μM) as single drugs and in the combinations shown, in triplicate experiments. Apoptosis was analyzed by Annexin V staining, measured by flow cytometry. *p=0.01; **p=0.008; ****p<.0001. **D.** NRG mice injected intravenously with M14(R)701-luc cells were treated with gilteritinib 7.5 mg/kg and/or FTY720 10 mg/kg, or vehicle control, starting on Day 1; serial non-invasive luciferin imaging of mice is shown. **E.** Changes in photon intensity, measured by bioluminescence imaging, over time, with *p*=0.015, comparing gilteritinib and FTY720 combination versus gilteritinib alone by 2-way ANOVA with Sidak’s multiple comparison test. **F.** Survival curves, with *p*=0.0035 comparing gilteritinib and FTY720 combination versus gilteritinib alone by Kaplan–Meier analysis.

M14(R)701 cells were treated with 10 nM gilteritinib or 1 nM quizartinib alone or in combination with 2.5 µM FTY720 or 5 µM KRP-203, another SIPR modulator, for 72 hours, and apoptosis was analyzed by Annexin V staining, measured by flow cytometry. Both FTY720 and KRP-203 in combination with gilteritinib or quizartinib markedly increased apoptosis, relative to single drugs (Figure 1C).

### FTY720 sensitizes M14(R)701 cells to gilteritinib *in vivo*

NRG mice injected intravenously with M14(R)701 cells dual-labelled with luciferase and GFP were treated with gilteritinib and/or FTY720 or vehicle control, as described in Methods. In this *in vivo* model, gilteritinib and FTY720 alone had little effect on luminescence, compared to vehicle control, but luminescence decreased significantly over time in mice treated with gilteritinib and FTY720 combination, compared with gilteritinib alone (p=0.015) (Figure 1D,E). Similarly, gilteritinib and FTY720 monotherapy modestly prolonged survival, compared to vehicle control, while gilteritinib and FTY720 combination significantly prolonged survival compared with gilteritinib alone (p=0.0035) (Figure 1F).

### FTY720 and quizartinib co-treatment inactivates ERK and AKT in M14(R)701 cells

To determine the effects of S1PR modulator and FLT3 inhibitor co-treatment on FLT3-ITD signaling, phosphorylated and total STAT5, ERK and AKT(S473) were measured in MOLM-14 cells and *NRAS* G12D-mutated M14(R)701 cells cultured with 1 nM quizartinib and/or 2.5 µM FTY720, or DMSO control, for two hours, compared to pre-treatment (Figure 2). p-STAT5, p-ERK and p-AKT(S473) decreased in MOLM-14 cells treated with quizartinib or quizartinib and FTY720, while p-STAT5, p-ERK and p-AKT did not decrease in M14(R)701 cells treated with quizartinib alone, but p-ERK and p-AKT decreased markedly following quizartinib and FTY720 co-treatment. In contrast, p-STAT5 did not decrease following quizartinib and FTY720 co-treatment.

**Figure 2.**
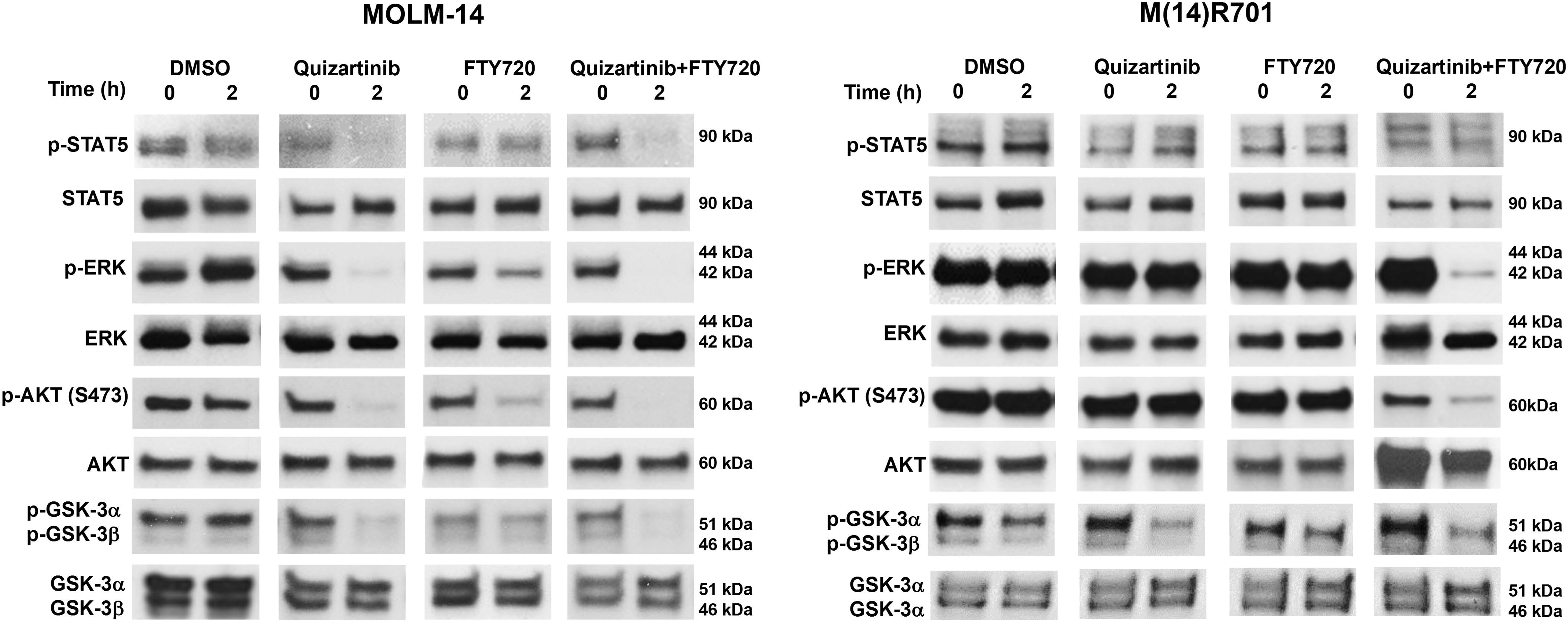
FTY720 and quizartinib co-treatment inactivates ERK and AKT in M14(R)701 cells. MOLM14 and M14(R)701 cells were treated with 1 nM quizartinib and/or 2.5 uM FTY720, or DMSO control, for 2 hours. p-STAT5, STAT, p-ERK, ERK, p-AKT, AKT, p-GSK-3α/β and GSK-3 α/β protein levels were measured by immunoblotting. Similar results were found in at least two independent experiments.

We previously reported FTY720 sensitization of FLT3-ITD AML cells to FLT3 inhibitors through activation of GSK-3β (18). Here we found that quizartinib and FTY720 co-treatment activates GSK-3β in MOLM-14 cells, consistent with our prior results, but less in M14(R)701 cells (Figure 2). Thus FTY720 does not sensitize M14(R)701 cells to FLT3 inhibitor through activation of GSK-3β.

### FLT3 inhibitor does not resensitize M14(R)701 cells through PP2A activation

The tumor suppressor PP2A is inactivated in FLT3-ITD AML cells (22,23), and co-treatment with PADs enhances FLT3 inhibitor efficacy in FLT3-ITD AML cell lines and patient blasts (18,22,23). We previously showed that PADs, including DT-061 and FTY720, enhance FLT3 inhibitor efficacy through AKT inhibition-dependent GSK-3β activation, increasing proteasomal degradation of c-Myc and Pim-1 (18). Additionally, concomitant targeting of SPHK1 was shown to exert synergistic cytotoxicity with FLT3 inhibitors through activation of the PP2A-GSK-3β pathway and consequent inhibition of β-catenin activity in FLT3-ITD cells (24).

Lack of GSK-3β activation in *NRAS*-mutated M14(R)701 cells treated with FTY720 and quizartinib relative to quizartinib alone (Figure 2), suggested that FTY720 resensitization of *NRAS*-mutated FLT3-ITD cells did not occur through the same mechanism as resensitization of FLT3-ITD cells without *NRAS* mutations, and specifically not through PP2A activation. To test this hypothesis, we studied the effects of the PAD DT-061 in M14(R)701 cells.

M14(R)701 cells were cultured with gilteritinib or quizartinib at increasing concentrations with or without 12 µM DT-061 (18). In contrast to FTY720 (Figure 1A), DT-061 did not resensitize M14(R)701 cells to FLT3 inhibitors (Figure 3A) or produce synergistic cytotoxicity in combination with gilteritinib or quizartinib at different combinations of concentrations (Figure 3B). Additionally, DT-061 and FLT3 inhibitor co-treatment did not increase apoptosis in M(R)701 cells (Figure 3C).

**Figure 3.**
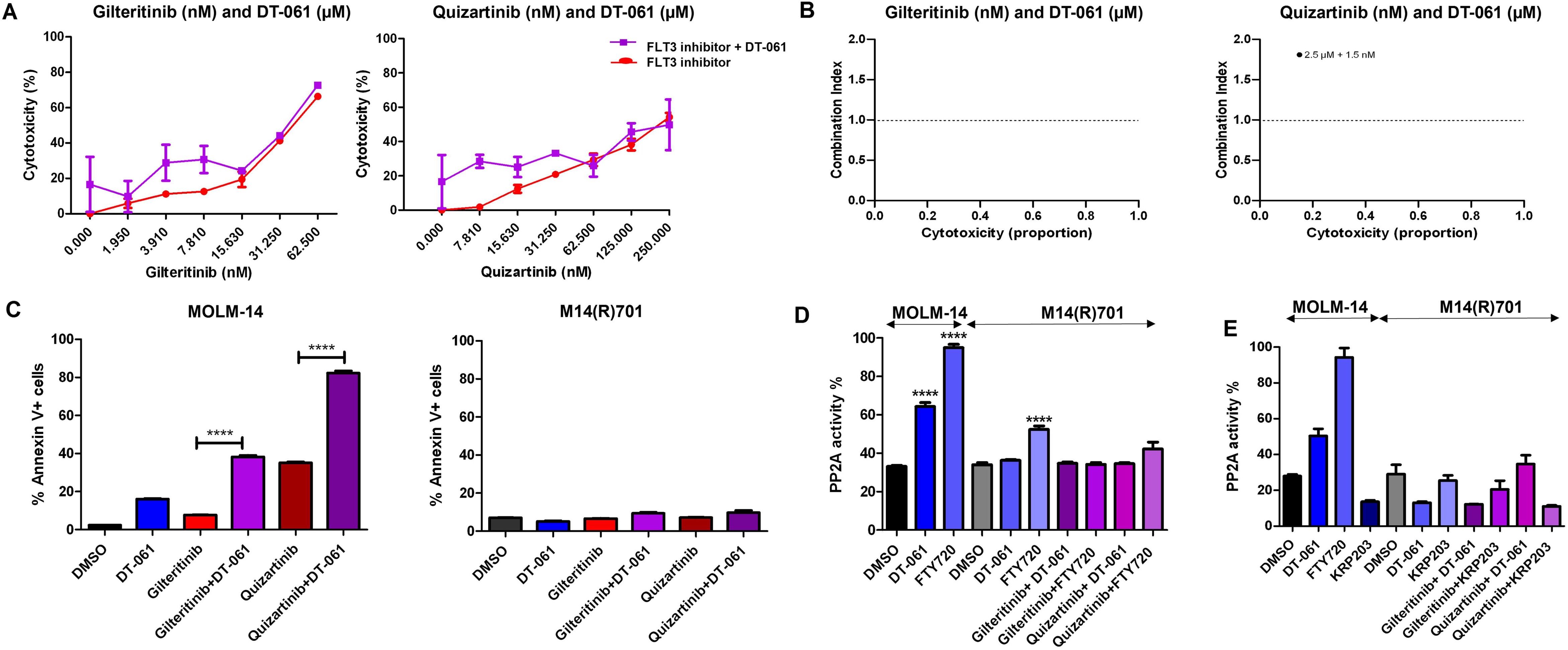
FLT3 inhibitor resensitization of M14(R)701 cells does not occur through PP2A activation. **A.** M14(R)701 cells were treated with the FLT3 inhibitors gilteritinib or quizartinib at increasing concentrations as single drugs and in combination with the PP2A-activating drug (PAD) DT-061 (12 μM) for 48 hours. Cytotoxicity was measured with the WST1 assay. **B.** M14(R)701 cells seeded at 10,000 cells/well in 96-well plates were treated for 48 hours with gilteritinib or quizartinib and/or DT-061 as single drugs and in combinations at diverse concentrations, in triplicate. Cytotoxicity was measured with the WST-1 assay, and drug combination effects were analyzed by the Chou-Talalay method. **C.** MOLM-14 and M14(R)701 cells were treated with the FLT3 inhibitors gilteritinib (10 nM) or quizartinib (1 nM) and/or the PAD DT-061 (12 μM) or DMSO control. Apoptosis was measured by Annexin V staining. **D.** PP2A phosphatase activity was measured in MOLM-14 and M14(R)701 cells treated with the PAD DT-061 (12 μM), the PAD and S1PR1 modulator FTY720 (2.5 μM) or the S1PR1 modulator KRP-203 (5 μM) and in M14(R)701 cells treated with the FLT3 inhibitors gilteritinib (10 nM) or quizartinib (1 nM) with or without DT-061, FTY720 or KRP-203 at the concentrations above. ****p<0.0001. Results are mean of two independent experiments.

To confirm that M14(R)701 cell resensitization to FLT3 inhibitors by FTY720, but not DT-061, was not due to differential efficacy of PP2A activation, PP2A phosphatase activity was measured in M14(R)701 and parental MOLM-14 cells treated with FTY720 or DT-061 alone and in combination with gilteritinib or quizartinib. Both FTY720 and DT-061 activated PP2A in parental MOLM14 cells, but FTY720 only modestly activated PP2A in M14(R)701 cells, and DT-061 did not (Figure 3D). Moreover, neither FTY720 nor DT-061 activated PP2A in combination with gilteritinib or quizartinib in M14(R)701 cells (Figure 3D). Similarly, KRP-203 did not increase PP2A phosphatase activity in parental MOLM-14 nor in *NRAS*-mutated M14(R)701 cells (Figure 3D), further demonstrating that S1PR modulator resensitization to FLT3 inhibitors does not occur through PP2A activation.

### S1PR modulator and FLT3 inhibitor co-treatment downregulates SPHK1

Given that SPHK1 is upregulated and localized to the plasma membrane in *RAS*-mutated cells (16,17), we studied effects of S1PR modulator and FLT3 inhibitor co-treatment on expression of SPHK1 in *NRAS*-mutated M14(R)710 cells.

FTY720 or KRP-203 and quizartinib co-treatment markedly downregulated SPHK1 protein in M14(R)701 cells, while FTY720 or KRP-203 alone had minimal or no effect (Figure 4A,B). SPHK1 protein turnover in M14(R)701 cells pretreated with CHX to block new protein translation prior to both combination treatments was similar to turnover with DMSO control Figure 4C,D). In contrast, SPHK1 mRNA measured by RT-PCR was downregulated in M14(R)701 cells treated with FTY720 and quizartinib for six hours (Figure 4E). Thus, SPHK1 downregulation by S1PR modulator and FLT3 inhibitor co-treatment in M14(R)701 cells was transcriptional.

**Figure 4.**
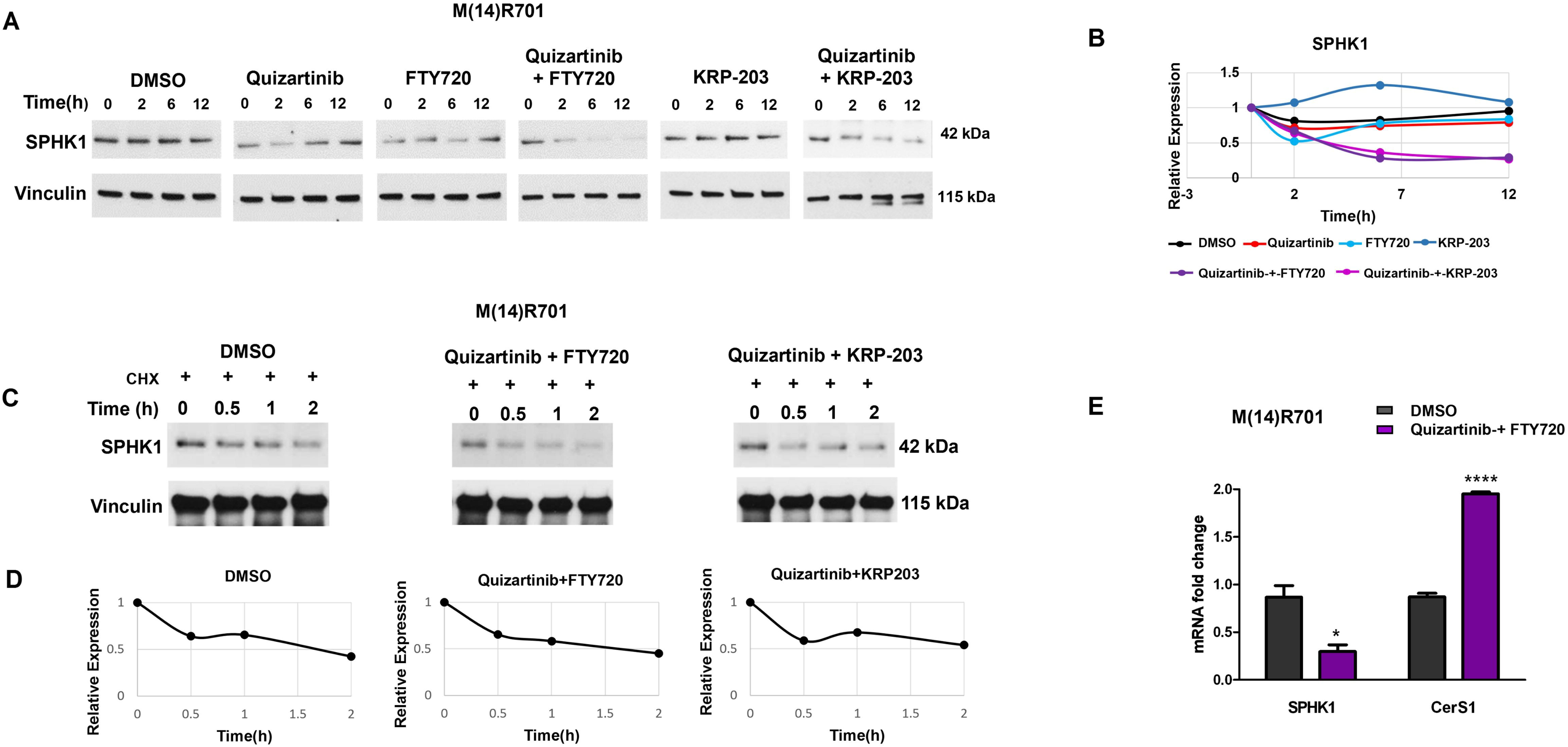
S1PR modulator and FLT3 inhibitor co-treatment downregulates SPHK1. **A.** M14(R)701 cells plated at 1.5x10^5^/ml were treated with quizartinib (1 nM) and/or FTY720 (2.5 μM) or KRP-203 (5 μM), or DMSO control. SPHK1 and vinculin loading control proteins were measured by immunoblotting in samples collected at serial time points. **B.** Densitometric analysis of data in **A** is shown, demonstrating downregulation of SPHK1 by FTY720 or KRP-203 in combination with quizartinib. Similar results were obtained in at least two independent experiments **C.** M14(R)701 cells plated at 1.5x10^5^/ml were treated with 100 µg/mL CHX for 1 hour to inhibit protein synthesis, before treatment with 1 nM quizartinib and/or 2.5 μM FTY720 or 5 μM KRP-203, or DMSO control. Samples collected at serial time points were analyzed for expression SPHK1 and vinculin loading control protein by immunoblotting. **D.** Densitometric analysis of data in **C** is shown, demonstrating that SPHK1 downregulation was not post-translational. Similar results were obtained in at least two independent experiments **E.** SPHK1 and ceramide synthase 1 (CerS1) mRNA expression was analyzed by RT-PCR in M14(R)701 cells before and after 6 hours of treatment with quizartinib (1 nM) and FTY720 (2.5 μM), or DMSO control. Data were normalized to 18S mRNA and graphed as fold change from Time 0 levels, defined as 1. SPHK1mRNA decreased, while CerS1 mRNA increased. Results are mean of two independent experiments. *p=0.01 ****p<0.0001

Downregulation of SPHK1 was accompanied by upregulation of ceramide synthase 1 (CerS1) mRNA (Figure 4E), indicating a shift in the sphingosine rheostat (25) from S1P synthesis toward ceramide synthesis.

### S1PR modulator and FLT3 inhibitor co-treatment inactivates AKT, p70 S6K and BAD

To examine the effects of S1PR modulator and FLT3 inhibitor combination treatment on signaling pathways downstream of *NRAS* (26), M14(R)701 cells treated with S1PR modulator and/or FLT3 inhibitor, or DMSO control, were harvested at serial time points and immunoblotted for phosphoproteins and total proteins.

Both FTY720 and quizartinib (Figure 5A,B) and gilteritinib and KRP-203 (Figure 5C,D) co-treatment rapidly and markedly downregulated p-AKT(S473) and (T308), but not total AKT, indicating AKT inactivation. Phosphorylation, but not total expression, of two downstream targets, p70 S6K and BAD, was also markedly downregulated by both co-treatments (Figure 5A-D).

**Figure 5.**
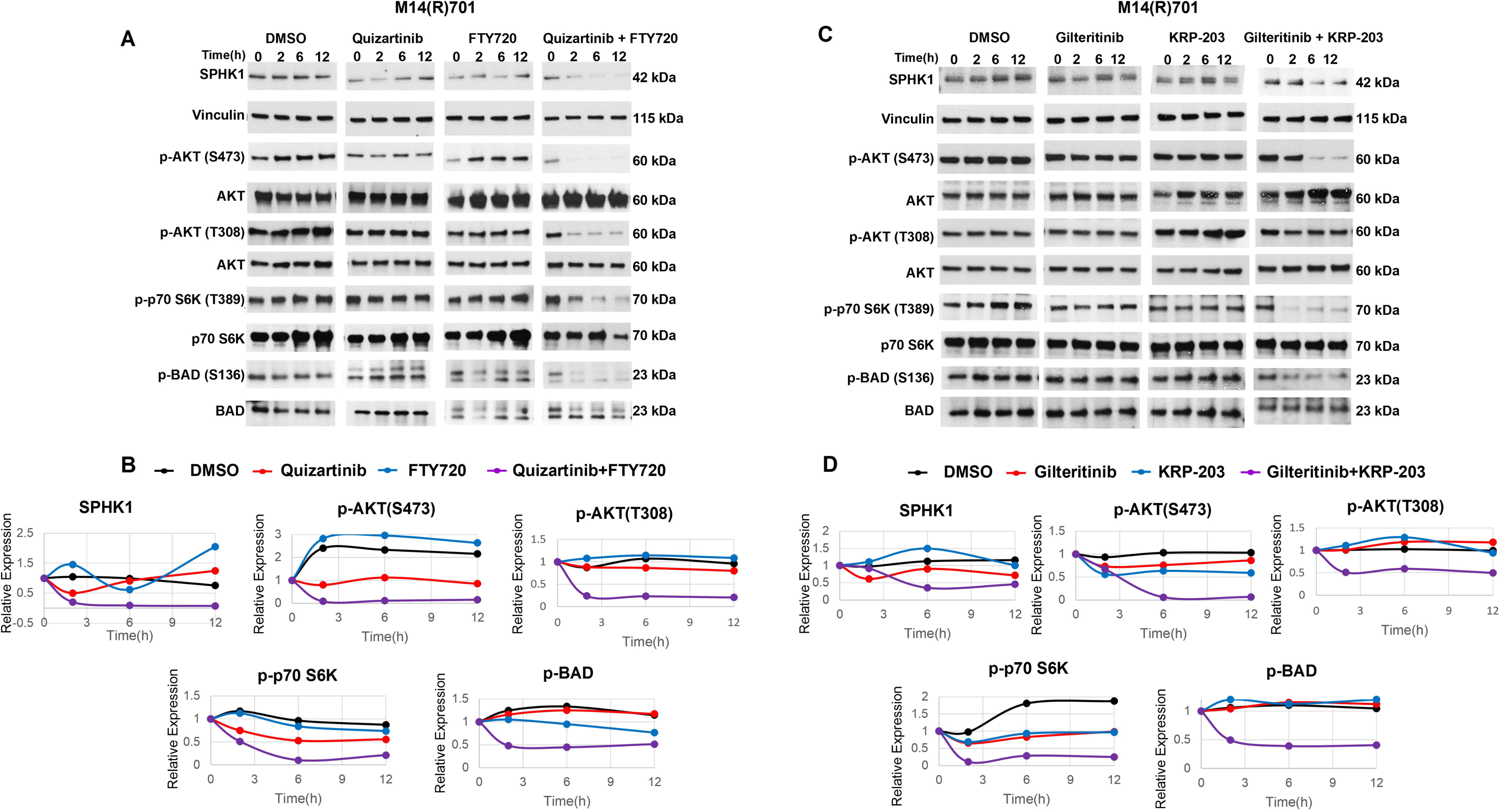
S1PR modulators resensitize M14(R)701 cells to FLT3 inhibitors via SPHK1/Akt/p70S6K/BAD pathway inhibition. **A.** M14(R)701 cells plated at 1.5 x10^5^/ml were treated with 1 nM quizartinib and/or 2.5 μM FTY720, or DMSO control. SPHK1, p-AKT/AKT, p-p70S6K/p70S6K, p-BAD/BAD and vinculin loading control proteins were measured at serial time points by immunoblotting. **B**. Densitometric analysis of data in **A** is shown, demonstrating downregulation of SPHK1, p-AKT, p-p70S6K and p-BAD by FTY720 and quizartinib combination treatment. Data quantified by densitometry and normalized to vinculin and Time 0 are shown graphically. **C.** M14(R)701 cells plated at 1.5 x10^5^/ml were treated with 10 nM gilteritinib and/or 5 μM KRP-203, or DMSO control. SPHK1, p-AKT/AKT, p-p70S6K/p70S6K, p-BAD/BAD and vinculin loading control proteins were measured at serial time points by immunoblotting. **D.** Densitometric analysis of data in **C** is shown, demonstrating downregulation of SPHK1, p-AKT, p-p70S6K and p-BAD by FTY720 and quizartinib combination treatment. Data quantified by densitometry and normalized to vinculin and Time 0 are shown graphically. Similar results were obtained in at least two independent experiments

### SPHK1 overexpression abrogates effects of S1PR modulator and FLT3 inhibitor co-treatment in M14(R)701 cells

To confirm that resensitization of M14(R)701 cells to FLT3 inhibitors by S1PR modulators is mediated through SPHK1, SPHK1 was overexpressed using a pLenti-SPHK1 construct, with a pLenti-empty vector as a control. Treatment of SPHK1-overexpressing M14(R)701 cells with 10 nM gilteritinib and 2.5 µM FTY720, did not downregulate SPHK1, as expected, and also did not downregulate p-AKT, p-p70S6K or p-BAD, while all four were downregulated in cells transduced with empty vector (Figure 6A,B). Furthermore, combination treatment induced apoptosis in cells transduced with empty vector, but not in SPHK1-overexpressing cells (Figure 6C). These findings support the conclusion that SPHK1 plays a critical role in mediating FLT3 inhibitor resensitization by S1PR modulators.

**Figure 6.**
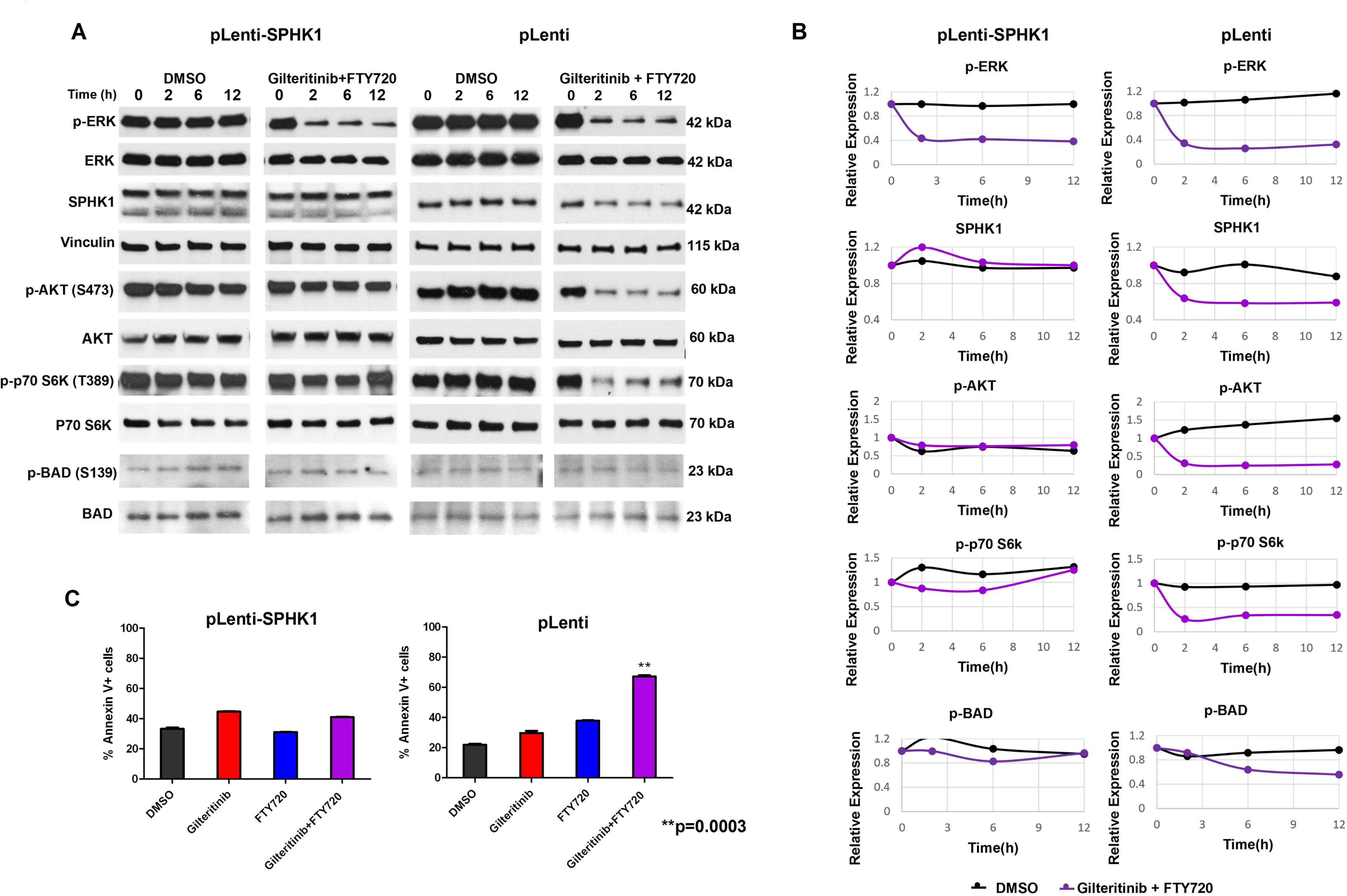
SPHK1 overexpression abrogates S1PR modulator resensitization of M14(R)701 cells to FLT3 inhibitor. **A.** M14(R)701 cells infected with pLenti SPHK1 plasmid or pLenti-empty vector were treated with 10 nM gilteritinib and/or 2.5 μM FTY720, or DMSO control. p-ERK/ERK, SPHK1, p-AKT/AKT, p-p70S6K/p70S6K, p-BAD/BAD and vinculin loading control protein levels were measured at serial time points by immunoblotting. **B.** Graphic representation of protein expression shown in **A** quantified by densitometry and normalized to vinculin and Time 0. Similar results were obtained in at least two independent experiments **C.** Apoptosis was measured by Annexin V staining in M14(R)701 cells infected with pLenti SPHK1 plasmid or pLenti-empty vector and treated with 10 nM gilteritinib and/or 2.5 μM FTY720, or DMSO control, for 72 hours. Apoptosis induction by gilteritinib and FTY720 combination was significantly reduced in cells infected with pLenti-SPHK1, compared with empty vector control. ***p=0.0003. Results are mean of two independent experiments.

Additionally, gilteritinib and FTY720 co-treatment downregulated p-ERK in M14(R)701 cells transduced with pLenti-SPHK1 and with empty vector (Figure 6A,B), confirming that ERK is upstream of SPHK1.

### Effect of S1PR modulator and FLT3 inhibitor co-treatment on FLT3-ITD cells with diverse *NRAS* mutations

We evaluated the effects of S1PR modulator and FLT3 inhibitor co-treatment in four additional NRAS-mutant cell lines: quizartinib-resistant MOLM-14(QS)-NRAS-G12C and MOLM-14(QS)-NRAS-Q61K and gilteritinib-resistant MOLM14-FGF2-G12S and MV411-FLT3-Q61H, with G12C, Q61K, G12S, and Q61H mutations, respectively. Apoptosis was measured by flow cytometry in all four cell lines treated with 10 nM gilteritinib and/or 2.5µM FTY720, or DMSO control, for 72 hours (Figure 7A). Combination treatment induced apoptosis in MOLM14-FGF2-G12S, MOLM-14(QS)-NRAS-Q61K and MV411-FLT3-Q61H cells with *NRAS* G12S, Q61K and Q61H mutations, respectively, but not in MOLM-14(QS)-NRAS-G12C cells, with an *NRAS* G12C mutation.

**Figure 7.**
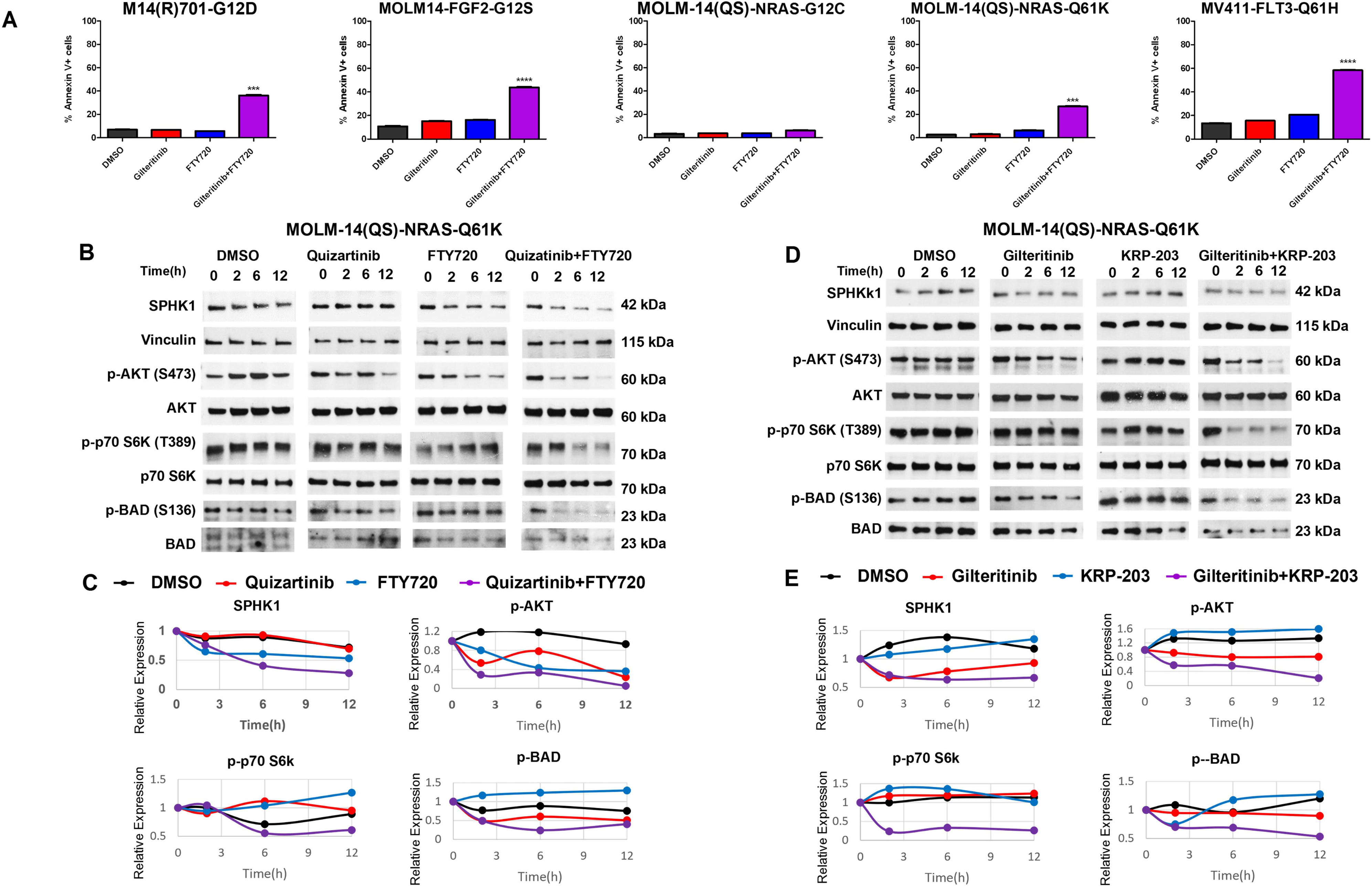
S1PR modulator and FLT3 inhibitor co-treatment of FLT3-ITD cells with diverse *NRAS* mutations. **A.** MOLM-14 and MV4-11 cell lines harboring the different *NRAS* mutations shown in the figure were treated with 10 nM gilteritinib and 2.5 μM FTY720. Apoptosis was measured by Annexin V staining. ***p=0.0002, ****p<0.0001. **B.** MOLM-14(QS)-NRASQ61K cells plated at 1.5 x10^5^/ml were treated with 1 nM quizartinib and/or 2.5 μM FTY720, or DMSO control. SPHK1, p-AKT/AKT, p-p70S6K/p70S6K, p-BAD/BAD and vinculin loading control proteins were measured at serial time points by immunoblotting**. C.** Data in **B** quantified by densitometry and normalized to vinculin and Time 0 are shown graphically. **D.** SPHK1, p-AKT/AKT, p-p70S6K/p70S6K, p-BAD/BAD and vinculin loading control proteins were measured at serial time points by immunoblotting in MOLM-14(QS)-NRASQ61K cells plated at 1.5 x10^5^/ml and treated with 10 nM gilteritinib and/or 5 μM KRP-203, or DMSO control. **E**. Data in **D** quantified by densitometry and normalized to vinculin and Time 0 are shown graphically. Similar results were obtained in at least two independent experiments

Quizartinib and FTY720 or gilteritinib and KRP-203 co-treatment downregulated SPHK1, p-AKT, p-p70 S6K, and p-BAD in MOLM-14(QS)-NRAS-Q61K cells with *NRAS* Q61K mutation (Figure 7B-E), similarly to M14(R)701 cells.

### Effect of S1PR modulator and FLT3 inhibitor co-treatment on FLT3-ITD AML patient blasts with *NRAS* mutations

Blasts from Patient 1, with FLT3-ITD and *NRAS* G13D, were treated with 1 nM quizartinib and/or 2.5 µM FTY720, or DMSO control, and harvested for immunoblotting at serial time points. SPHK1, p-AKT, p-p70 S6K and p-BAD were downregulated by combination treatment, but not by single-drug treatments (Figure 8A,B). Additionally, cytotoxicity was measured after 48 hours with the WST-1 assay. The combination treatment produced markedly greater cytotoxicity, compared to single drugs (Figure 8C). Blasts from Patient 2, with G13V, were treated for 6 hours with 1 nM quizartinib and/or 2.5 µM FTY720, or DMSO control, and harvested for immunoblotting. SPHK1, p-AKT, p-p70 S6K and p-BAD were downregulated by combination treatment, but not by single-drug treatments (Figure 8D,E).

**Figure 8.**
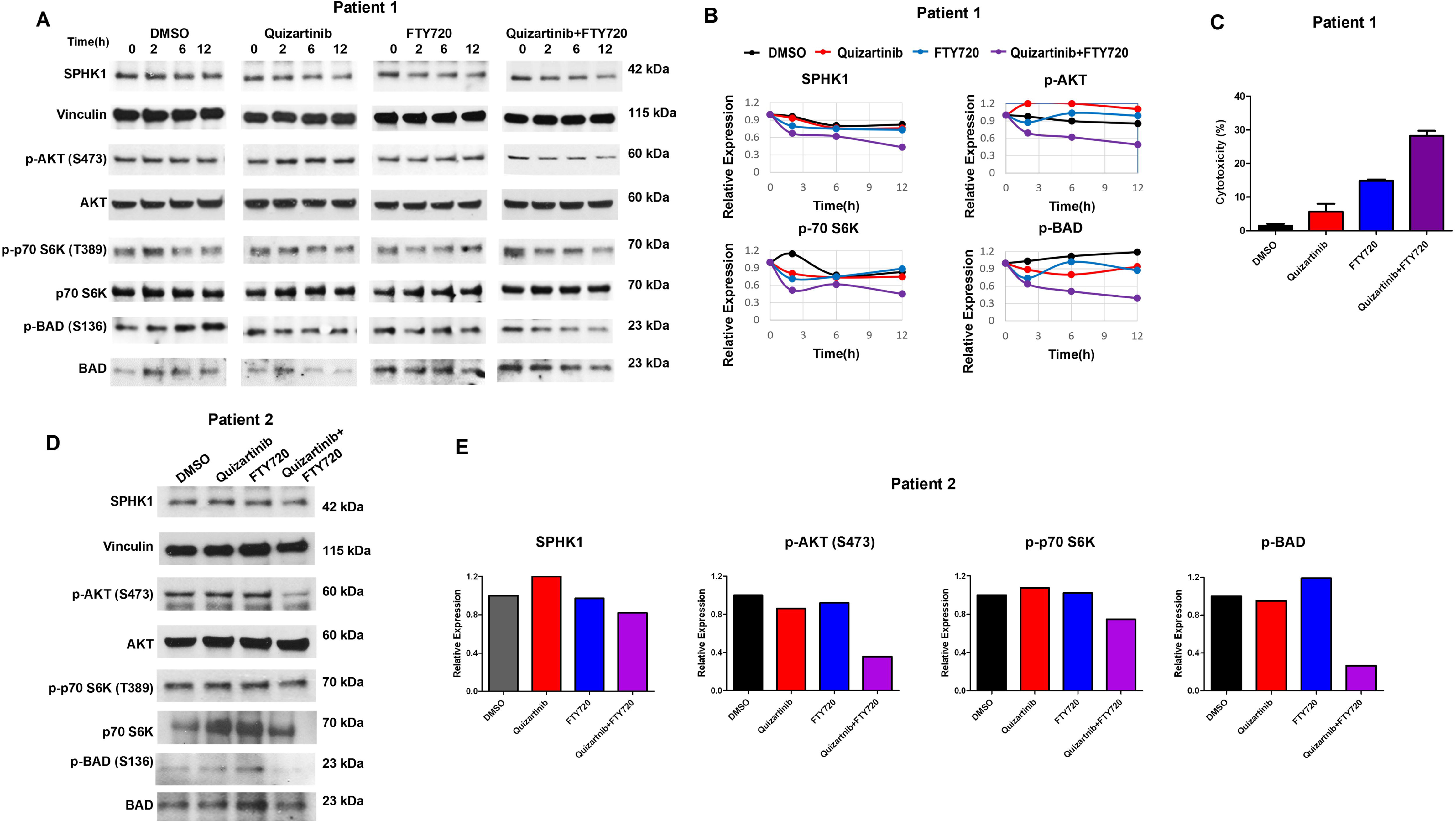
S1PR modulator and FLT3 inhibitor co-treatment of AML patient blasts with FLT3-ITD and G13D or G13V *NRAS* mutations. **A.** Blasts from Patient 1, with FLT3-ITD and an *NRAS* G13D mutation treated with 1nM quizartinib and/or 2.5µM FTY720, or DMSO control, were harvested for immunoblotting at serial time points and SPHK1, p-AKT/AKT, p-p70 S6K/p70 S6K and p-BAD/BAD protein expression was measured. **B**. Graphic representation of data in **A** is shown. SPHK1, p-AKT, p-p70 S6K and p-BAD were downregulated by combination treatment, but not by single-drug treatments. **C.** Blasts from Patient 1 were incubated for 48 hours with 1nM quizartinib and/or 2.5µM FTY720, or DMSO control, and cytotoxicity was measured using the WST-1 assay. Combination treatment produced markedly greater cytotoxicity, compared to single drugs. **D.** Blasts from Patient 2, with FLT3-ITD and an *NRAS* G13V mutation, were treated with 1nM quizartinib and/or 2.5µM FTY720, or DMSO control, for 6 hours, and cells were harvested for immunoblotting. **E.** Graphic representation of the data in **D** is shown. SPHK1, p-AKT/AKT, p-p70 S6K/p70 S6K and p-BAD/BAD were downregulated by combination treatment, but not by single-drug treatments.

## Discussion

FLT3-ITD, present in AML cells in 25% of patients, is associated with hyperleukocytosis, high relapse rate, rapid relapses and poor patient outcomes. Incorporation of FLT3 tyrosine kinase inhibitors into treatment of FLT3-ITD AML has improved outcomes (4–6), but responses are not durable (5), with onset of resistance frequently associated with development of new mutations (7). *NRAS* is the most commonly mutated gene in FLT3-ITD AML with acquired FLT3 inhibitor resistance (9–13), with point mutations altering NRAS protein codons G12, G13 or Q61 constitutively activating RAS proteins by impairing GTPase activity, causing constitutive Ras-Raf-MEK-ERK signaling (14). Here we found that co-treatment with S1PR modulators resensitizes FLT3-ITD-expressing cells with G12D, G12S, Q61K and Q61H, G13D and G13V, albeit not G12C, *NRAS* mutations to FLT3 inhibitors. The S1PR modulator fingolimod (FTY720) is FDA-approved for treatment of multiple sclerosis and has been considered safe for treatment of AML (27). Three other S1PR modulators, siponimod, ozanimod, and ponesimod, are also approved for treatment of multiple sclerosis (28). The S1PR modulator mocravimod (KRP-203) has been safely used in clinical trials for prevention of graft-versus-host disease in patients with hematologic malignancies undergoing allogeneic hematopoietic stem cell transplantation (29). Therefore, the combination treatments studied here have the potential to be developed clinically.

FLT3-ITD signals constitutively through PI3K-AKT-mTOR and Ras-Raf-MEK-ERK, as well as, aberrantly, though STAT5 (3). We found that S1PR modulator and FLT3 inhibitor combination treatment inactivates ERK and AKT, but not STAT5, in *NRAS* G12D-mutated M14(R)701 cells. Thus, the combination treatment inactivates constitutive Ras-Raf-MEK-ERK signaling downstream of activated *NRAS*. Constitutive ERK activation is a well-recognized mechanism of FLT3 inhibitor resistance, not only downstream of mutated *RAS*, but also due to cues from the bone marrow stroma (30) as well as intrinsic mechanisms (31).

Activating *NRAS* mutations sustain aberrant ERK signaling, which promotes SPHK1 expression and activity, linking oncogenic RAS to sphingolipid metabolism. Increased SPHK1 expression shifts the balance of the sphingolipid rheostat toward S1P, activating AKT and downstream p70S6K and BAD, while suppressing ceramide-driven apoptosis, thus contributing to survival and drug resistance of AML cells. The findings position SPHK1 as a critical effector integrating the RAS/ERK and AKT survival pathways.

FTY720 sensitizes FLT3-ITD cells with and without *NRAS* mutations to FLT3 inhibitors by different mechanisms. We previously showed that the PADs FTY720 and DT-061 enhance sensitivity of FLT3-ITD cells to FLT3 inhibitors through activation (dephosphorylation) of GSK-3β and consequent enhanced proteasomal degradation of c-Myc and Pim-1 (18). We show here that quizartinib and FTY720 combination treatment activates GSK-3β in MOLM-14 cells, but not in *NRAS*-mutated M14(R)701 cells. We further show that both FTY720 and DT-061, another PAD, activate PP2A in MOLM-14 cells, but not in M14(R)701 cells.

Sphingosine has been implicated in mutant *RAS* signaling. SPHK1 activity was significantly increased in NIH3T3 cells transfected with activated mutant H-Ras (V12-Ras), compared to wild-type *RAS,* and treatment with a specific SPHK1 inhibitor or co-transfection with inactive SPHK1 reduced proliferation of mutant RAS-transfected cells (32). HEK293T cells transfected with G12V mutant K-Ras showed increased S1P production and decreased ceramide production, mediated by increased SPHK1 activity (33). Moreover, constitutively active B-Raf or MEK1 activated SPHK1, while constitutively active Akt1 did not, indicating that increased S1P production is mediated by the Raf/MEK/ERK pathway (33). SPHK1 is also overexpressed downstream of ERK activation in melanoma cells with BRAF or NRAS mutations (*NRAS* Q61L or *BRAF* V600E), compared with normal melanocytes (34).

The sphingosine rheostat has also been implicated in FLT3-ITD signaling and FLT3 inhibitor efficacy and resistance. Pro-apoptotic lipid ceramide levels were lower in cells with FLT3-ITD, compared to wild-type FLT3, and FLT3 inhibition in FLT3-ITD cells increased synthesis and mitochondrial accumulation of ceramide, causing lethal mitophagy (35). Treatment with the mitochondria-targeted ceramide analog LCL-461 resensitized resistant MV4-11 cells to the FLT3 inhibitor crenolanib, though the mechanism of crenolanib resistance was not specified (35). Moreover, long-term sorafenib treatment upregulated SPHK1/S1P signaling in FLT3-ITD cells, and SPHK1 inhibition potently enhanced FLT3 inhibitor efficacy *in vitro* and *in vivo* (36). Mechanistically, targeting SPHK1/S1P/S1PR2 signaling synergized with FLT3 inhibitors to activate the PP2A-GSK-3β pathway and thereby inhibit β-catenin activity (36). In contrast, we show here that the efficacy of co-targeting SPHK1/S1P and FLT3 signaling is not due to PP2A-GSK-3β pathway activation in *NRAS*-mutated cells. Finally, co-inhibition of FLT3 and SPHK1 resensitized midostaurin-resistant FLT3-ITD-expressing MV4-11 and MOLM-13 cells, inducing apoptosis, though the mechanism of midostaurin resistance was not characterized (37). We show here that quizartinib and FTY720 or KRP-203 co-treatment of FLT3-ITD AML cells with most *NRAS* mutations modulates sphingosine signaling, with SPHK1 downregulation and CerS1 upregulation, indicating a shift of sphingosine signaling from S1P to ceramide.

Here, co-treatment with S1PR modulators resensitized FLT3-ITD-expressing cells with G12D, G12S, Q61K and Q61H, G13D and G13V, but not G12C, *NRAS* mutations to FLT3 inhibitors. Early work on RAS inhibition focused on KRAS, which is commonly mutated in solid tumors (38). Two KRAS inhibitors, sotorasib and adagrasib, were developed (39,40), with efficacy limited to G12C *KRAS* mutations, as inhibition depends on covalent binding with mutant cysteine 12, locking KRAS in the inactive state (41,42). Sotorasib has some activity against NRAS G12C, while adagrasib does not (43). In our work here, S1PR modulator and FLT3 inhibitor co-treatment was effective against FLT3-ITD AML cells with all *NRAS* mutations studied except G12C, possibly due to the biochemical properties of the cysteine substitution at codon 12, which introduces a reactive thiol group altering NRAS conformation, membrane localization, and effector interactions, possibly modifying effects on downstream signaling modules, including SPHK1.

Broader RAS inhibitors have been developed, including the pan-KRAS inhibitors BI-2493 and BI-2865 (44) and the pan-RAS inhibitors RMC-7977 (45), RMC-6236 (46) and ADT-007 (47), which inhibit KRAS, NRAS and HRAS with diverse mutations as well as activated wild-type RAS. The pan-RAS inhibitor RMC-7977 resensitizes FLT3-ITD AML cell lines and patient samples with *RAS* mutations to FLT3 inhibitors *in vitro* and *in vivo*, with ERK inactivation and variable AKT inhibition, likely downstream of ERK, as with S1PR modulators and FLT3 inhibitors here (48).

Results here support a unified model in which activated RAS/ERK-driven SPHK1 signaling sustains AKT activation and supports a pro-survival lipid state. Pharmacologic modulation of this pathway may offer a promising approach to overcoming FLT3 inhibitor resistance in FLT3-ITD AML with *NRAS* mutations and potentially other mutations activating Ras-Raf-MEK-ERK signaling.

## Acknowledgements

The authors gratefully acknowledge the patients who donated samples under a University of Maryland School of Medicine IRB-approved tissue procurement protocol, Lamis Farah, B.S., and colleagues, University of Maryland Greenebaum Comprehensive Cancer Center, for patient sample procurement, the University of Maryland Greenebaum Comprehensive Cancer Genomics Shared Resource for molecular analyses, and Pathology and Biorepository Shared Resource and Translational Laboratory Shared Resource for tissue banking.

## Author Contributions

MRB, AC, MKMA and GS conceptualized the study and designed the experiments. AC and GS performed and analyzed the *in vitro* experiments. CMB and YW performed the xenograft experiment. DS, CCS and ET created and shared *NRAS*-mutated resistant cell lines. YL contributed intellectually. AC and MRB wrote the manuscript. All authors reviewed, contributed to and approved the final version of the manuscript.

## Data Availability Statement

All relevant data are included in the article and the supplementary file. Data are available from the corresponding author upon reasonable request.

## Graphical Abstract

Summary figure showing the effects of concurrent treatment with S1PR modulators and FLT3 inhibitors in FLT3-ITD AML cells with *NRAS* mutations.

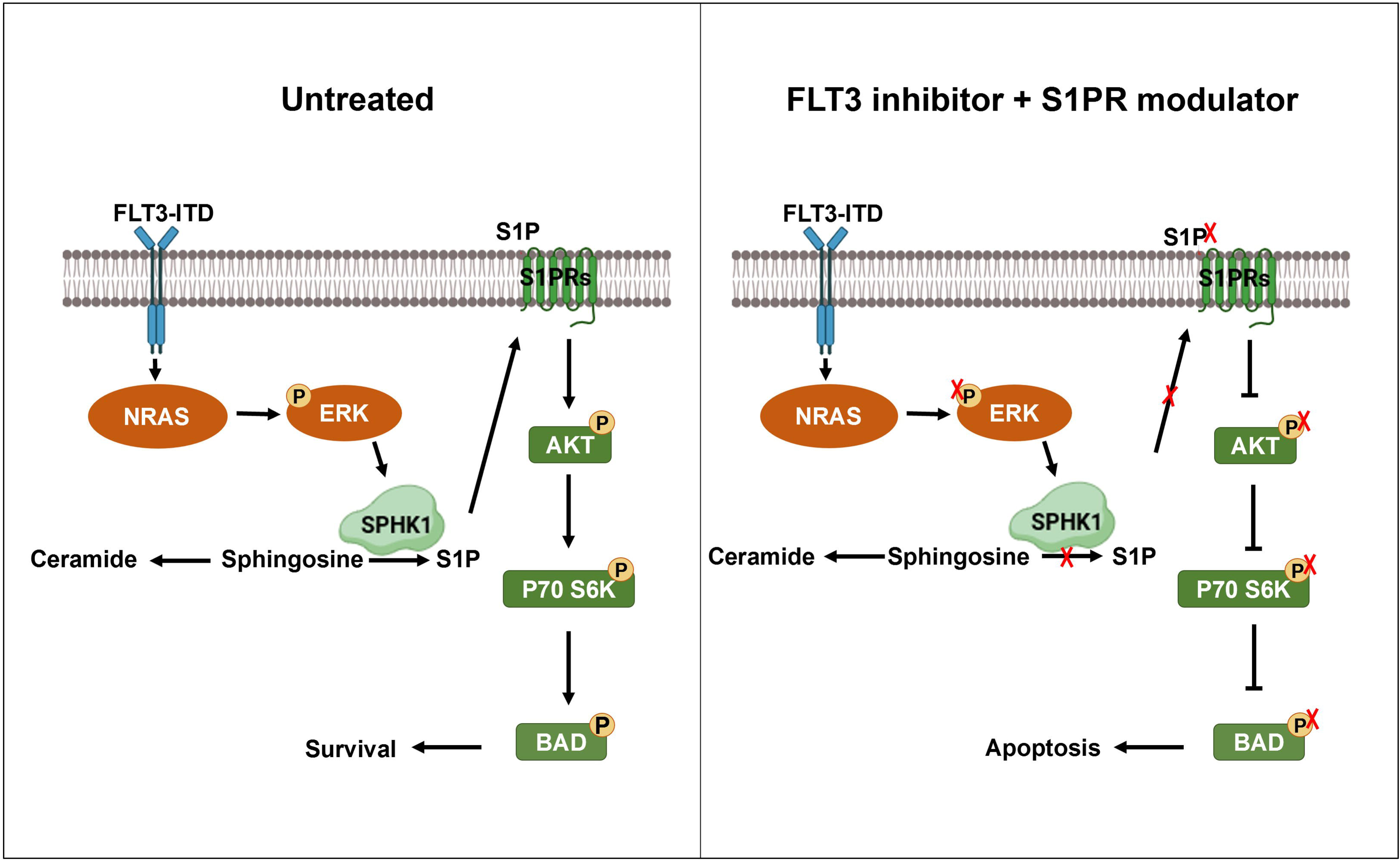

## Supplementary

**Supplementary Table S1.** Patient samples.

| Patient | Age, sex | AML status | WBC (x10<sup>9</sup>/L) | Blasts (%) | Karyotype | *FLT3*-ITD size (bp) | Insertion length | *FLT3*-ITD allelic burden (%) | *NRAS* mutation | *NRAS* mutation VAF (%) | Other mutated genes |
| --- | --- | --- | --- | --- | --- | --- | --- | --- | --- | --- | --- |
| 1 | 58M | Diagnosis | 92 | 78 | 46,XY | 376 | 47 | 38 | G13D | 12 | *NPM1* |
| 2 | 53F | Relapse 1 | 174 | 80 | 46,XX, t(6;9)(p22;q34) | 405 | 77 | 111 | G13V | 49 | *WT1* |

**Supplementary Table S2.**
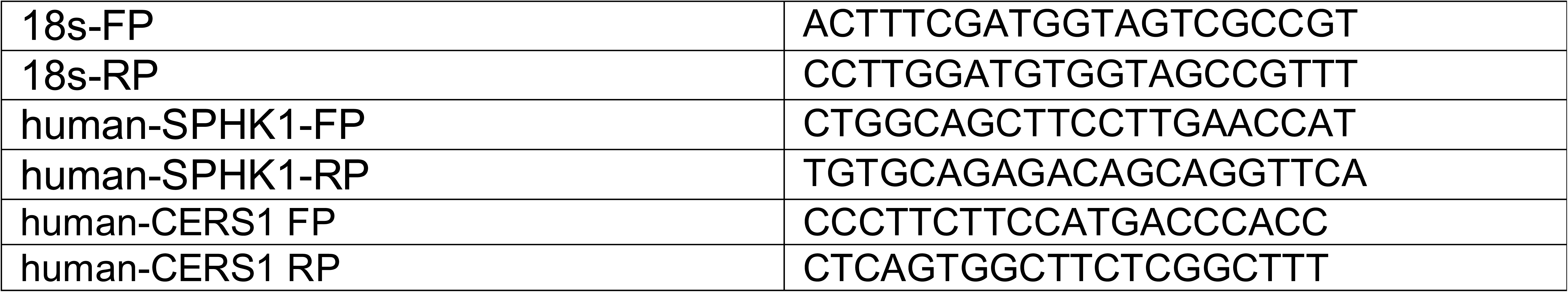
Primers used for qRT-PCR.

**Supplementary Table S3.**
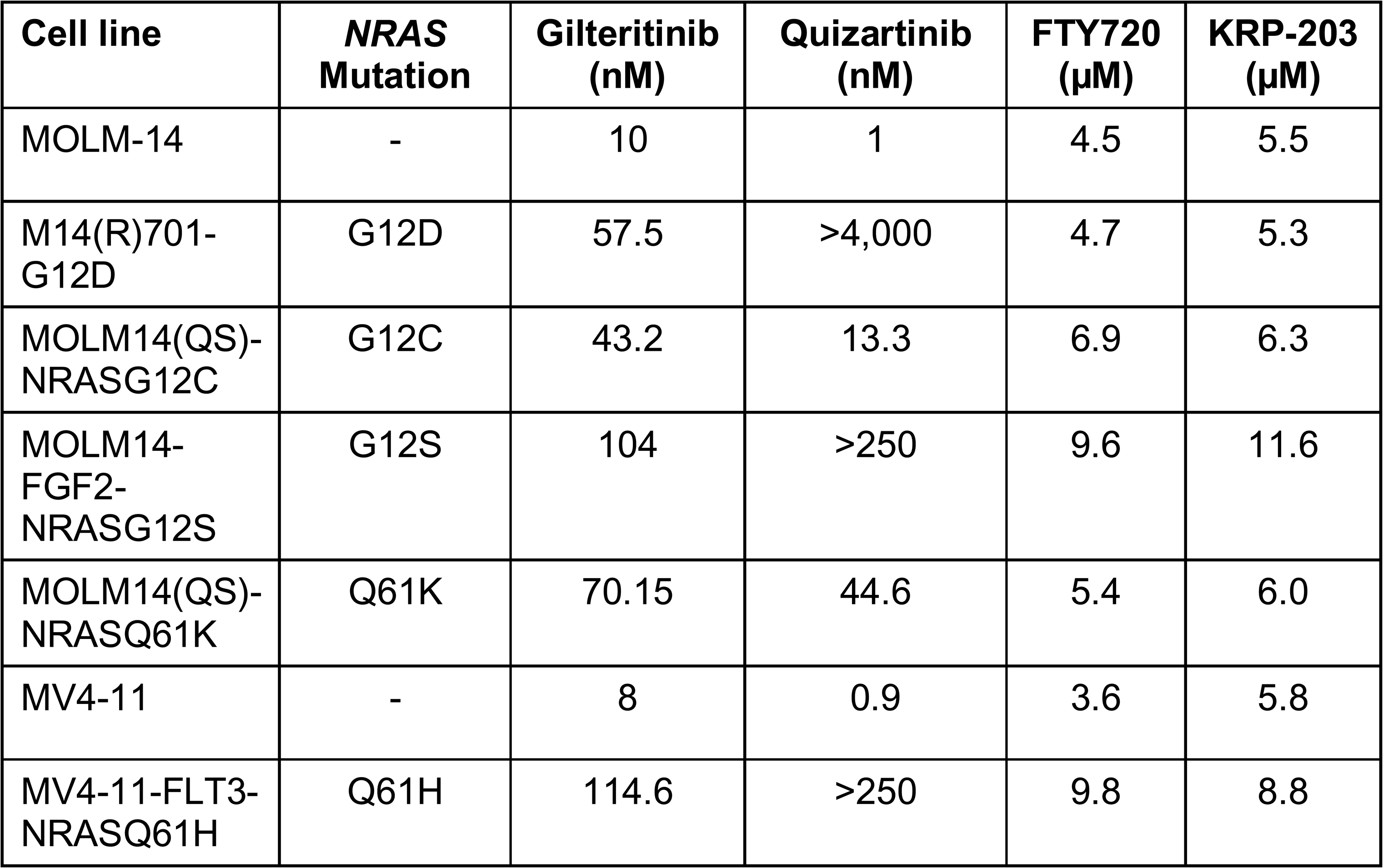
Drug IC_50_s in parental and *NRAS*-mutated FLT3-ITD AML cell lines.

## References

1. Patel JP, Gönen M, Figueroa ME, Fernandez H, Sun Z, Racevskis J, et al. Prognostic relevance of integrated genetic profiling in acute myeloid leukemia. N Engl J Med. 2012;366:1079–1089.

2. Papaemmanuil E, Gerstung M, Bullinger L, Gaidzik VI, Paschka P, Roberts ND, et al. Genomic Classification and Prognosis in Acute Myeloid Leukemia. N Engl J Med. 2016;374:2209–2221.

3. Spiekermann K, Bagrintseva K, Schwab R, Schmieja K, Hiddemann W. Overexpression and constitutive activation of FLT3 induces STAT5 activation in primary acute myeloid leukemia blast cells. Clin Cancer Res. 2003;9:2140–2150.

4. Stone RM, Mandrekar SJ, Sanford BL, Laumann K, Geyer S, Bloomfield CD, et al. Midostaurin plus Chemotherapy for Acute Myeloid Leukemia with a FLT3 Mutation. N Engl J Med. 2017;377:454–464.

5. Perl AE, Martinelli G, Cortes JE, Neubauer A, Berman E, Paolini S, et al. Gilteritinib or Chemotherapy for Relapsed or Refractory *FLT3*-Mutated AML. N Engl J Med. 2019;381:1728–1740.

6. Erba HP, Montesinos P, Kim HJ, Patkowska E, Vrhovac R, Žák P, et al. Quizartinib plus chemotherapy in newly diagnosed patients with FLT3-internal-tandem-duplication-positive acute myeloid leukaemia (QuANTUM-First): a randomised, double-blind, placebo-controlled, phase 3 trial. Lancet. 2023;401:1571–1583.

7. Kennedy VE, Smith CC. FLT3 targeting in the modern era: from clonal selection to combination therapies. Int J Hematol. 2024;120:528–540.

8. Piloto O, Wright M, Brown P, Kim KT, Levis M, Small D. Prolonged exposure to FLT3 inhibitors leads to resistance via activation of parallel signaling pathways. Blood. 2007;109:1643–1652.

9. Zhang H, Savage S, Schultz AR, Bottomly D, White L, Segerdell E, et al. Clinical resistance to crenolanib in acute myeloid leukemia due to diverse molecular mechanisms. Nat Commun. 2019;10:244.

10. McMahon CM, Ferng T, Canaani J, Wang ES, Morrissette JJD, Eastburn DJ, et al. Clonal Selection with RAS pathway activation mediates secondary clinical resistance to selective FLT3 inhibition in acute myeloid leukemia. Cancer Discov. 2019;9:1050–1063.

11. Alotaibi AS, Yilmaz M, Kanagal-Shamanna R, Loghavi S, Kadia TM, DiNardo CD, et al. Patterns of resistance differ in patients with acute myeloid leukemia treated with type I versus type II FLT3 inhibitors. Blood Cancer Discov. 2021;2:125–134.

12. Smith CC, Levis MJ, Perl AE, Hill JE, Rosales M, Bahceci E. Molecular profile of FLT3-mutated relapsed/refractory patients with AML in the phase 3 ADMIRAL study of gilteritinib. Blood Adv. 2022;6:2144–2155.

13. Joshi SK, Nechiporuk T, Bottomly D, Piehowski PD, Reisz JA, Pittsenbarger J, et al. The AML microenvironment catalyzes a stepwise evolution to gilteritinib resistance. Cancer Cell. 2021;39:999.e8–1014.e8.

14. Prior IA, Lewis PD, Mattos C. A comprehensive survey of Ras mutations in cancer. Cancer Res. 2012;72:2457–2467.

15. Larrosa-Garcia M, Baer MR. FLT3 inhibitors in acute myeloid leukemia: Current status and future directions. Mol Cancer Ther. 2017;16:991–1001.

16. Gault CR, Eblen ST, Neumann CA, Hannun YA, Obeid LM. Oncogenic K-Ras regulates bioactive sphingolipids in a sphingosine kinase 1-dependent manner. J Biol Chem. 2012;287:31794–31803.

17. Zhu W, Gliddon BL, Jarman KE, Moretti PAB, Tin T, Parise LV, et al. CIB1 contributes to oncogenic signalling by Ras via modulating the subcellular localisation of sphingosine kinase 1. Oncogene. 2017;36:2619–2627.

18. Scarpa M, Singh P, Bailey CM, Lee JK, Kapoor S, Lapidus RG, et al. PP2A-activating drugs enhance FLT3 inhibitor efficacy through AKT inhibition-dependent GSK-3β-mediated c-Myc and Pim-1 proteasomal degradation. Mol Cancer Ther. 2021;20:676–690.

19. Lee JK, Chatterjee A, Scarpa M, Bailey CM, Niyongere S, Singh P, et al. Pim kinase inhibitors increase gilteritinib cytotoxicity in FLT3-ITD acute myeloid leukemia through GSK-3β activation and c-Myc and Mcl-1 proteasomal degradation. Cancer Res Commun. 2024;4:431–445.

20. Chou TC. Drug combination studies and their synergy quantification using the Chou-Talalay method. Cancer Res. 2010;70:440–446.

21. Neviani P, Santhanam R, Oaks JJ, Eiring AM, Notari M, Blaser BW, et al. FTY720, a new alternative for treating blast crisis chronic myelogenous leukemia and Philadelphia chromosome-positive acute lymphocytic leukemia. J Clin Invest. 2007;117:2408–2421.

22. Agarwal A, MacKenzie RJ, Pippa R, Eide CA, Oddo J, Tyner JW, et al. Antagonism of SET using OP449 enhances the efficacy of tyrosine kinase inhibitors and overcomes drug resistance in myeloid leukemia. Clin Cancer Res. 2014;20:2092–2103.

23. Smith AM, Dun MD, Lee EM, Harrison C, Kahl R, Flanagan H, et al. Activation of protein phosphatase 2A in FLT3+ acute myeloid leukemia cells enhances the cytotoxicity of FLT3 tyrosine kinase inhibitors. Oncotarget. 2016;7:47465–47478.

24. Jiang L, Zhao Y, Liu F, Huang Y, Zhang Y, Yuan B, et al. Concomitant targeting of FLT3 and SPHK1 exerts synergistic cytotoxicity in FLT3-ITD^+^ acute myeloid leukemia by inhibiting β-catenin activity via the PP2A-GSK3β axis. Cell Commun Signal. 2024;22:391.

25. Newton J, Lima S, Maceyka M, Spiegel S. Revisiting the sphingolipid rheostat: Evolving concepts in cancer therapy. Exp Cell Res. 2015;333:195–200.

26. Cox AD, Der CJ. “Undruggable KRAS”: druggable after all. Genes Dev. 2025; 39:132–162.

27. Enjeti AK, D’Crus A, Melville K, Verrills NM, Rowlings P. A systematic evaluation of the safety and toxicity of fingolimod for its potential use in the treatment of acute myeloid leukaemia. Anticancer Drugs. 2016;27:560–568.

28. McGinley MP, Cohen JA. Sphingosine 1-phosphate receptor modulators in multiple sclerosis and other conditions. Lancet. 2021;398:1184–1194.

29. Dertschnig S, Gergely P, Finke J, Schanz U, Holler E, Holtick U, et al. Mocravimod, a selective sphingosine-1-phosphate receptor modulator, in allogeneic hematopoietic stem cell transplantation for malignancy. Transplant Cell Ther. 2023;29:41.e1–e9.

30. Yang X, Sexauer A, Levis M. Bone marrow stroma-mediated resistance to FLT3 inhibitors in FLT3-ITD AML is mediated by persistent activation of extracellular regulated kinase. Br J Haematol. 2014;164:61–72.

31. Bruner JK, Ma HS, Li L, Qin ACR, Rudek MA, Jones RJ, et al. Adaptation to TKI treatment reactivates ERK signaling in tyrosine kinase-driven leukemias and other malignancies. Cancer Res. 2017;77:5554–5563.

32. Xia P, Gamble JR, Wang L, Pitson SM, Moretti PA, Wattenberg BW, et al. An oncogenic role of sphingosine kinase. Curr Biol. 2000;10:1527–1530.

33. Gault CR, Eblen ST, Neumann CA, Hannun YA, Obeid LM. Oncogenic K-Ras regulates bioactive sphingolipids in a sphingosine kinase 1-dependent manner. J Biol Chem. 2012;287:31794–31803.

34. Albinet V, Bats ML, Huwiler A, Rochaix P, Chevreau C, Ségui B, et al. Dual role of sphingosine kinase-1 in promoting the differentiation of dermal fibroblasts and the dissemination of melanoma cells. Oncogene. 2014;33:3364–3373.

35. Dany M, Gencer S, Nganga R, Thomas RJ, Oleinik N, Baron KD, et al. Targeting FLT3-ITD signaling mediates ceramide-dependent mitophagy and attenuates drug resistance in AML. Blood. 2016;128:1944–1958.

36. Jiang L, Zhao Y, Liu F, Huang Y, Zhang Y, Yuan B, et al. Concomitant targeting of FLT3 and SPHK1 exerts synergistic cytotoxicity in FLT3-ITD^+^ acute myeloid leukemia by inhibiting β-catenin activity via the PP2A-GSK3β axis. Cell Commun Signal. 2024;22:391.

37. Tecik M, Adan A. Concurrent inhibition of FLT3 and sphingosine kinase-1 triggers synergistic cytotoxicity in midostaurin resistant FLT3-ITD positive acute myeloid leukemia cells via blocking FLT3/STAT5A signaling to induce apoptosis. J Chemother. 2025:1–17.

38. Simanshu DK, Nissley DV, McCormick F. RAS Proteins and Their Regulators in Human Disease. Cell. 2017;170:17–33.

39. Hong DS, Fakih MG, Strickler JH, Desai J, Durm GA, Shapiro GI, et al. KRAS^G12C^ inhibition with sotorasib in advanced solid tumors. N Engl J Med. 2020;383:1207–1217.

40. Ou SI, Jänne PA, Leal TA, Rybkin II, Sabari JK, Barve MA, et al. First-in-human phase I/IB dose-finding study of adagrasib (MRTX849) in patients with advanced *KRAS*^G12C^ solid tumors (KRYSTAL-1). J Clin Oncol. 2022;40:2530–2538.

41. Ostrem JM, Peters U, Sos ML, Wells JA, Shokat KM. K-Ras(G12C) inhibitors allosterically control GTP affinity and effector interactions. Nature. 2013;503:548–551.

42. Lito P, Solomon M, Li LS, Hansen R, Rosen N. Allele-specific inhibitors inactivate mutant KRAS G12C by a trapping mechanism. Science. 2016;351:604–608.

43. Rubinson DA, Tanaka N, Fece de la Cruz F, Kapner KS, Rosenthal MH, Norden BL, et al. Sotorasib is a pan-RASG12C inhibitor capable of driving clinical response in NRASG12C cancers. Cancer Discov. 2024;14:727–736.

44. Tedeschi A, Schischlik F, Rocchetti F, Popow J, Ebner F, Gerlach D, et al. Pan-KRAS inhibitors BI-2493 and BI-2865 display potent antitumor activity in tumors with KRAS wild-type allele amplification. Mol Cancer Ther. 2025;24:550–562.

45. Holderfield M, Lee BJ, Jiang J, Tomlinson A, Seamon KJ, Mira A, et al. Concurrent inhibition of oncogenic and wild-type RAS-GTP for cancer therapy. Nature. 2024;629:919–926.

46. Jiang J, Jiang L, Maldonato BJ, Wang Y, Holderfield M, Aronchik I, et al. Translational and Therapeutic Evaluation of RAS-GTP Inhibition by RMC-6236 in RAS-Driven Cancers. Cancer Discov. 2024;14:994–1017.

47. Foote JB, Mattox TE, Keeton AB, Chen X, Smith FT, Berry K, et al. A pan-RAS inhibitor with a unique mechanism of action blocks tumor growth and induces antitumor immunity in gastrointestinal cancer. Cancer Res. 2025;85:956–972.

48. Popescu B, Jones MF, Piao M, Tran E, Koh A, Lomeli I, et al. Multiselective RAS(ON) inhibition targets oncogenic RAS and overcomes RAS-mediated resistance to FLT3i and BCL2i in AML. Blood. 2025 Oct 27:blood.2025030558. doi: 10.1182/blood.2025030558. Epub ahead of print.

